# Inference of gene regulatory networks for overcoming low performance in real-world data

**DOI:** 10.1101/2024.07.16.603684

**Authors:** Yusuke Hiki, Yuta Tokuoka, Takahiro G. Yamada, Akira Funahashi

**Affiliations:** Center for Biosciences and Informatics, Graduate School of Fundamental Science and Technology, Keio University, Yokohama, Kanagawa, 223-8522, Japan; Department of Biosciences and Informatics, Keio University, Yokohama, Kanagawa, 223-8522, Japan

## Abstract

The identification of gene regulatory networks is important for understanding the mechanisms of various biological phenomena. Many methods have been proposed to infer networks from time-series gene expression data obtained by high-throughput next-generation sequencings. Such methods can effectively infer gene regulatory networks for *in silico* data, but inferring the networks accurately from *in vivo* data remiains a challenge because of the large noise and low time sampling rate. Here, we proposed a novel unsupervised learning method, Multi-view attention Long-short term memory for Network inference (MaLoN). It can infer gene regulatory networks with temporal changes in gene regulation using the multi-view attention Long Short-term memory model. Using *in vivo* benchmark datasets in *Saccharomyces cerevisiae* and *Escherichia coli*, we showed that MaLoN can infer gene regulatory networks more accurately than existing methods. The ablated models indicated that the multi-view attention mechanism suppressed false positives. The order of activation of gene regulations inferred by MaLoN was consistent with existing knowledge.

## Introduction

Gene regulation plays a key role in regulating mRNA and protein abundance and governs various biological phenomena such as cell cycle, cell fate determination, and responses to environmental stresses [1–3]. Identification of gene regulatory networks, which represent regulatory relationships among genes, makes it possible, for example, to identify drug targets [4] and master regulators in diseases [5]. The identification of such networks is one of the key challenges in systems biology.

Gene regulatory networks can be inferred with steady-state models or time-series models. The former type infers indirect causal relationships from samples in steady state; it can infer robust and accurate networks based on many samples under a variety of perturbation conditions, but cannot capture direct causal relationships among genes [6]. The latter can capture direct causal relationships from time-series samples under various perturbations, but usually only a limited number of samples (data points) is available before a steady state is reached.

Many methods based on the time-series model have been reported to highly accurately infer gene regulatory networks from a few data points [7–14]. A fast and simple method represents gene regulation by a linear function and the time derivative of target gene expression as a weighted linear sum of the expression of regulatory factors [7]. But gene regulation is generally represented by a nonlinear function, and then the regression of target gene expression cannot be sufficiently learned. Samples used to obtain time-series gene expression data are often sparse because of the high cost, and the approximation to calculate the time derivative using adjacent time points results in large errors. Therefore, the performance of modeling the time derivative strongly depends on the time sampling frequency.

Bayesian network models can directly infer network structures to explain time-series expression data [8–10]. They predict causal relationships among genes as the structure with the highest posterior probability; given an infinite number of data points, the posterior probabilities for gene regulation asymptotically approach the true probability distribution. These methods can also deal with noise, but an increase in the number of genes would lead to a combinatorial explosion and computational difficulties.

As a machine learning method, regression tree models enable nonlinear regression and are computationally feasible for many genes [11–14]. The differential expression of the target gene is detected and the regression tree is constructed, which regresses the expression from genes that best explain the difference as the variable for splitting. The gene for splitting that can reduce the regression error more is regarded as an important variable that contributes to the regression of the target gene and has a high score representing the plausibility of a regulatory relationship that explains temporal causality. A regression tree with each target gene is independently trained to infer scores for the regulatory relationship among all genes without any ground truth data on true gene regulations. Such trees are fast even for multivariate data and prevent overfitting by assembling weak regressors, thus enabling highly accurate inference even for noisy data.

GENIE3, an inference method using Random Forests (RF), a type of regression tree, is a steady-state model with the highest performance for the DREAM4 challenge dataset generated *in silico* as a benchmark [15]. An extension of GENIE3, dynGENIE3 can also be applied to timeseries data [12]. JUMP3 divides gene regulation into promoter regulation by transcription factors and transcription based on promoter activity; it models the former with a regression tree model using RF and the latter with RF with stochastic differential equations [11]. BiXGBoost [13] and NonlinearODEs [14] replace XGBoost [16] based on gradient boosting and allow for more flexible and scalable inference. BiXGBoost predicts both candidate regulators and candidate target genes from regression and then integrates the predictions. NonlinearODEs use XGBoost to regress the time derivative of the target gene expression.

However, the above methods only focus on local relationships such as time *t −* 1 and *t −* 2 to regress expression of genes at time *t*. *In silico* data are generated on basis of ordinary differential equations (ODE) and stochastic differential equations (SDE) models and then rely only on local temporal relationships. In *in vivo* data, the time lag in which temporal causality is observed is unknown, and depends on experimental conditions. The user must speculate on the number of timepoints as a lag for regression, which makes it impossible to track dynamically changing gene regulations [2, 17, 18]. Because the number of time series for *in vivo* data is often small, using expression patterns from earlier time points as background information would allow for more appropriate inference. Methods based on regression tree models are highly accurate for *in silico* data, but the performance for *in vivo* data is low [19].

Recurrent neural networks (RNNs) can directly learn the global temporal feature of time-series data by memorizing past time-series and perform nonlinear regression to represent gene regulation. However, RNNs are uninterpretable and have not been applied to gene regulatory network inference, which requires variable selection in regression. The attention mechanism is effective in making RNNs interpretable, which requires variable selection in regression [20, 21].

In this study, we developed an unsupervised learning method, Multiview attention Long-short term memory for Network inference (MaLoN) for highly accurate inference of gene regulatory networks from *in vivo* time-series gene expression data. MaLoN was applied to 3 *in vivo* datasets and outperformed the regression tree–based methods, which are currently the most accurate. We evaluated how the novel features of MaLoN, the multi-view attention mechanism and Long-Short Term Memory, contribute to its performance by comparing MaLoN with ablated models. The multi-view attention mechanism especially effectively suppressed false positives. We showed that MaLoN can predict several known dynamic gene regulations and therefore has the potential to reveal the timing of activation of gene regulations.

## Results

### Performance on the IRMA and *Escherichia coli* SOS pathway datasets

MaLoN, implemented in this study, is depicted in Figure 1. MaLoN consists of a Long-Short Term Memory (LSTM) model with a time- and gene-view attention mechanism to represent dynamic changes in gene regulation by extracting global features of time-series data. Our algorithm learns regressions of gene expression with an LSTM model that incorporates attention mechanisms for both gene and time-view and quantifies the plausibility scores, which reveals a regulatory relationship for each gene. We compared MaLoN with regression-based gene regulatory network inference algorithms that use machine learning: JUMP3 [11], BiXGBoost [13], dynGENIE3 [12], and NonlinearODEs [14]. We used the *in vivo* assessment of reverse-engineering and modeling assessment (IRMA; switch-on, switch-off) dataset, a time-series gene expression data based on a synthetic network in *S. cervisiae* regulated by galactose [22], and the *E. coli* SOS Pathway dataset, which is based on the response network to DNA damage [23]. These datasets are shown in Table 1. As the metrics for evaluation, we used area under the precision-recall curve (AUPR), area under the receiver operating characteristic curve (AUROC), and Max F-measure. AUPR and AUROC indicate how accurately the ranking of the inferred scores corresponds to the network of correct answers, and Max F-measure is the highest F-measure among the networks obtained by adjusting thresholds. Our method outperformed the existing methods in these metrics (Fig. 2): the average AUROC, AUPR, and Max F-measure on the three datasets improved by 0.019, 0.141, and 0.191. We visualized the gene regulatory network obtained using the threshold corresponding to the Max F-measure (Fig. 3, Supplementary Figs. 1 and 2). On average, MaLoN predicted the network with the fewest errors and the fewest false positives. Only MaLoN accurately inferred the substructures known to generate complex dynamics, such as the feedback loop [24] between the CBF1, SWI5, and GAL4 nodes in the IRMA switch-on dataset.

**Fig. 1:**
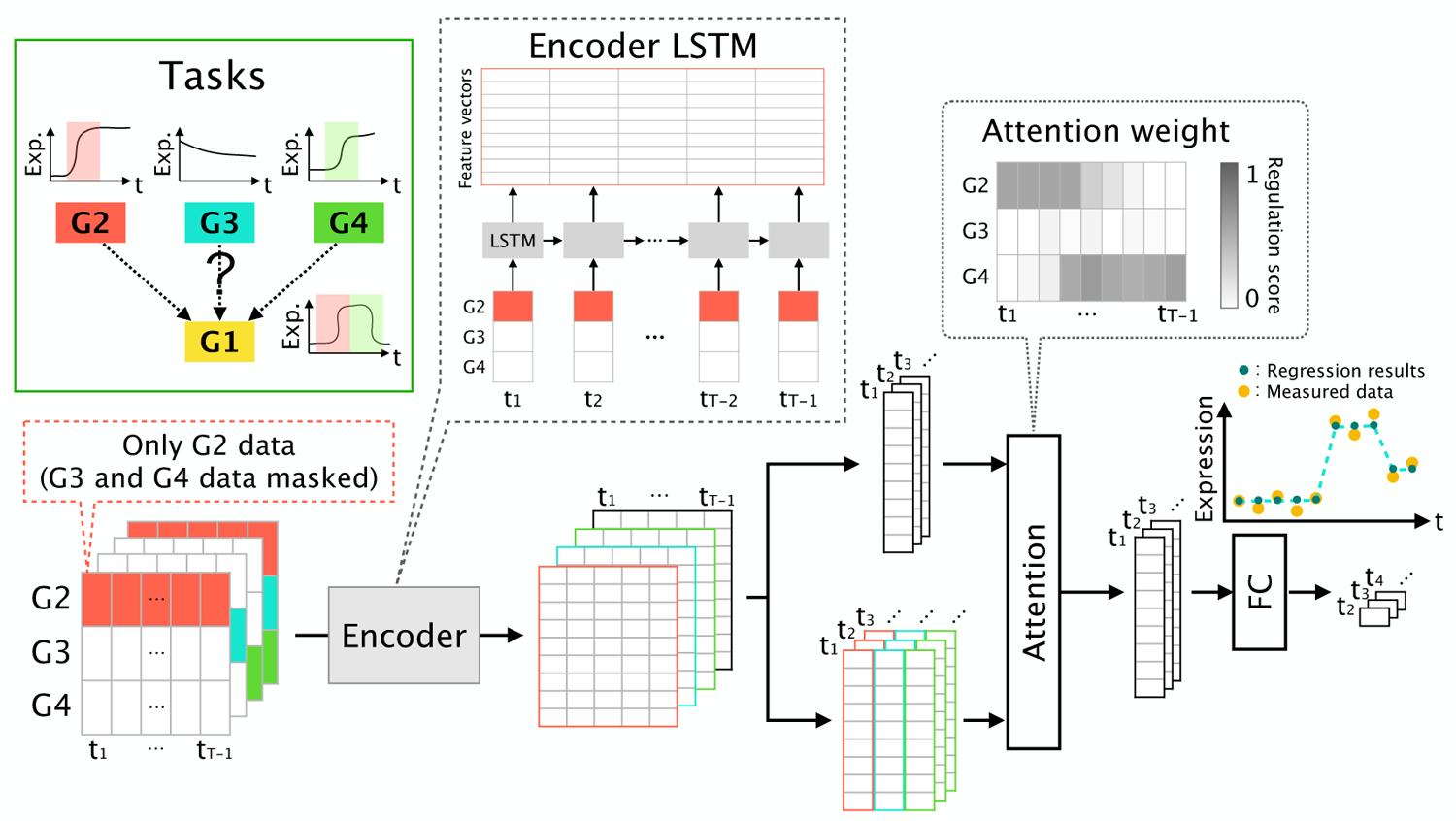
Overview of our MaLoN algorithm. The tasks are to train a model to regress the expression of target genes from the other genes and to quantify which genes are likely to regulate the target genes by interpreting the importance of the variables to the regression. First, Long-Short Term Memory (LSTM) models are used to obtain features from the time-series expression of each gene and all genes. Then the features are integrated using attention based on similarity for the features obtained from each gene and all genes. The integrated feature vector is used to regress the expression of target genes at the next time point. Attention weight is calculated as the similarity of the feature vector obtained from each gene and all genes.

**Fig. 2:**
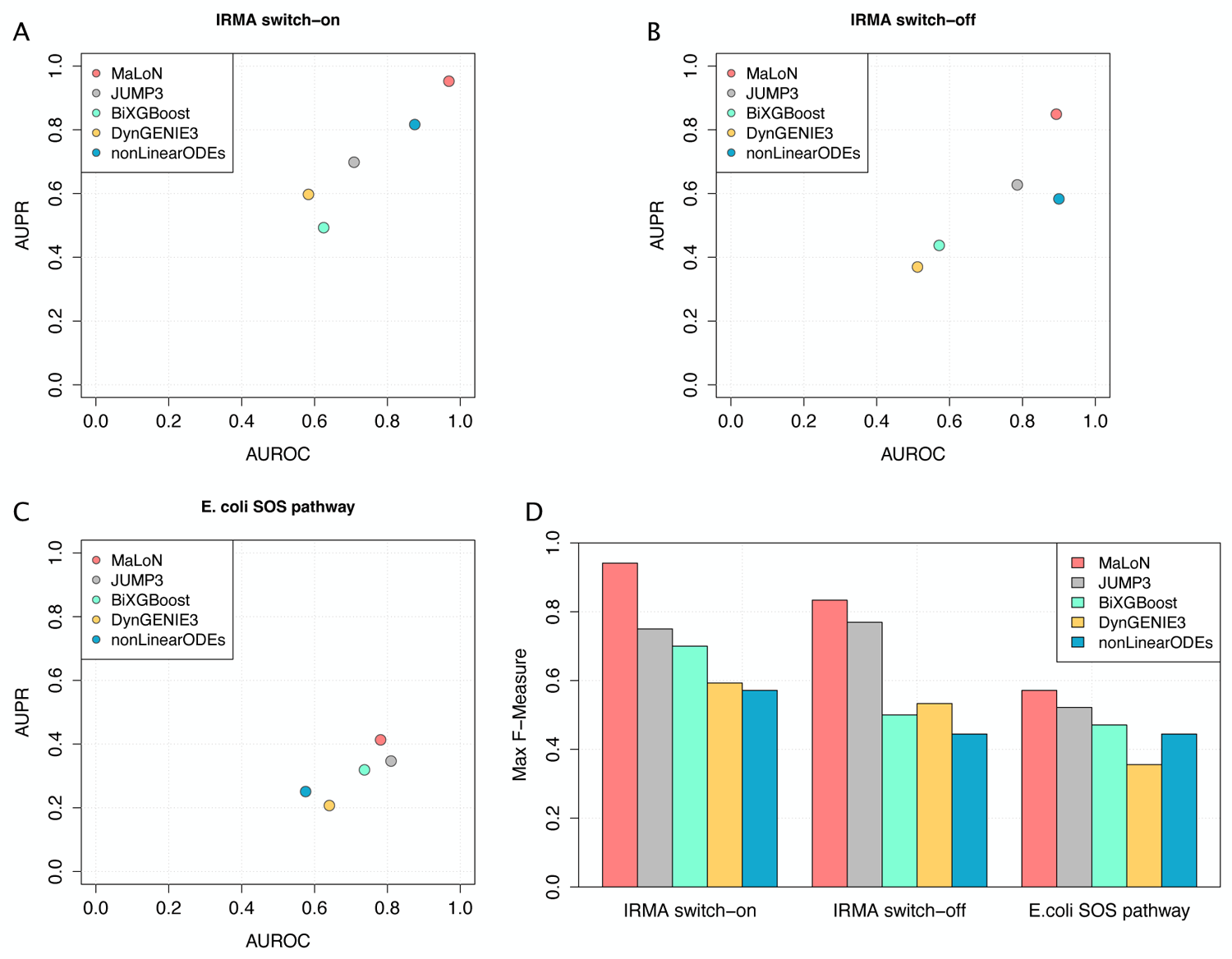
Comparison of MaLoN and other machine learning methods JUMP3 [11], BiXGBoost [13], DynGENIE3 [12], NonlinearODEs [14]. (a-c) AUROC, AUPR and (d) Max F-measure were used as metrics. Datasets used are indicated on the *x* axis in (d)

**Fig. 3:**
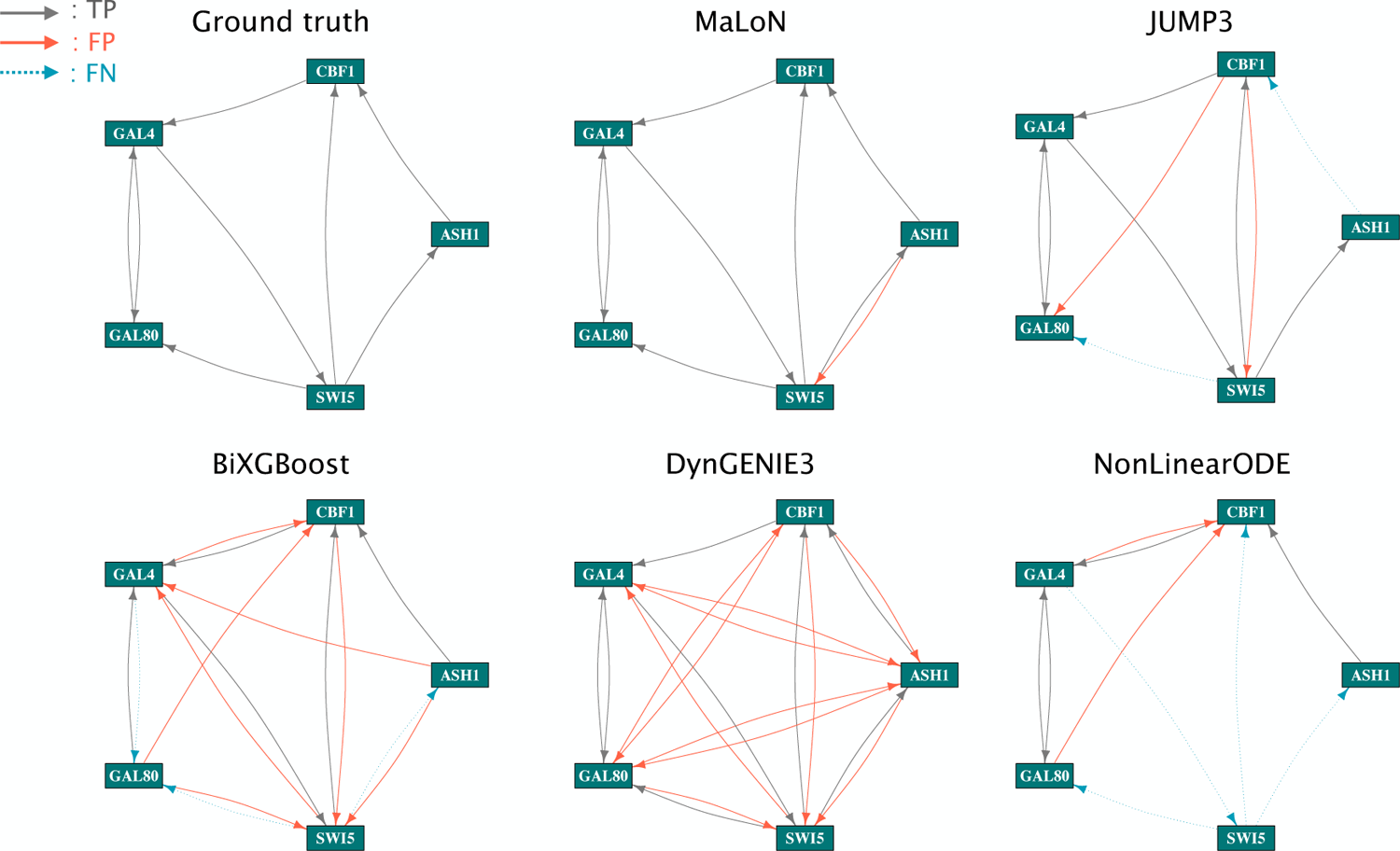
Visualization of inferred gene regulatory network in IRMA switch-on dataset. The nodes represent the 5 genes of this dataset, TP, true positive; FP, false positive; and FN, false negative. Ground truth is the true network, and the other networks were obtained by each method using the threshold with Max F-measure.

**Table 1:**
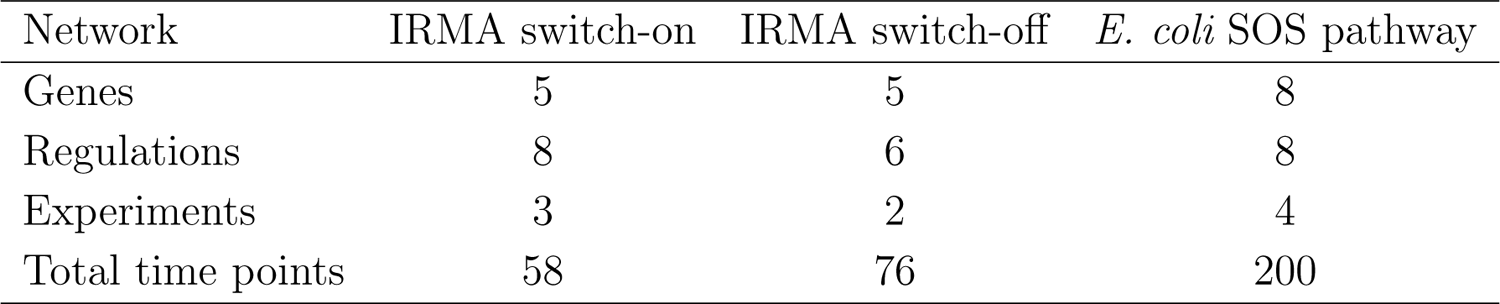
Datasets used in performance evaluation.

### Ablated models for Multi-view attention and LSTM

In MaLoN, we used LSTM incorporating multi-view attention mechanism (Mv-LSTM) for temporal causal detection among gene expression. To evaluate how the novel features (the multi-view attention mechanism and LSTM) contribute to the performance, we compared MaLoN with ablated models. We constructed single-view attention LSTM models (Sv-LSTM), in which the attention mechanism was applied only in the geneview (Fig. 4A), and the multi-view attention DNN (Mv-DNN), in which LSTM was replaced with a neural network with only fully connected layers (Fig. 4B). For the Mv-DNN architecture, we constructed two models: Mv-DNN10 and Mv-DNN36. In the former, the size of the feature vectors from the encoder was set to 10, as in Mv-LSTM (MaLoN), and to 36 in the latter, so that the number of model parameters was the same as in Mv-LSTM.

**Fig. 4:**
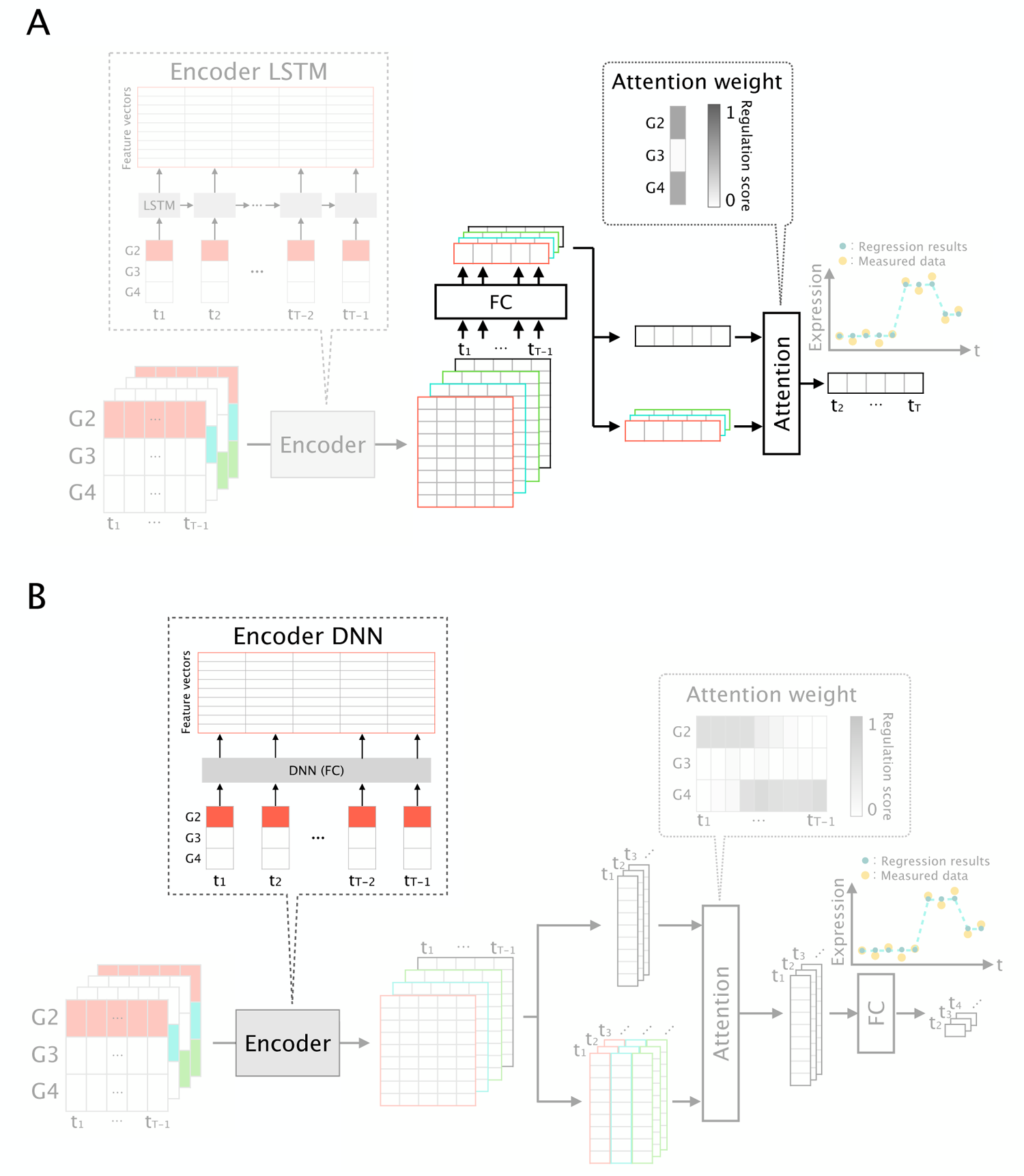
Overview of MaLoN-based ablated models for comparison with the original MaLoN. The different parts from MaLoN are highlighted. (A) Single-view attention LSTM model. To evaluate the contribution of multi-view attention (Mv), the attention layer was changed to singleview, which has gene-view. This model cannot represent dynamic gene regulation. (B) Multi-view attention Deep Neural Network (Mv-DNN). To evaluate the contribution of LSTM, it was replaced with a DNN consisting of fully connected layers. This model can extract local information but not global time-series features.

Mv-LSTM had the highest performance in AUROC, AUPR, and Max F-Measure (Fig. 5). Moreover, the difference in performance between Mv-LSTM and Sv-LSTM was the largest between Mv-LSTM and Sv-LSTM, indicating that the multi-view attention mechanism was especially important for accurate gene regulatory network inference. Precision and recall in the predicted network with Max F-Measure showed that Mv-LSTM greatly improved precision compared to the ablated models (Supplementary Table 1). The adoption of both multi-view attention mechanism and LSTM were effective in suppressing false positives, and in particular, multi-view attention mechanism contributed greatly to improving performance.

**Fig. 5:**
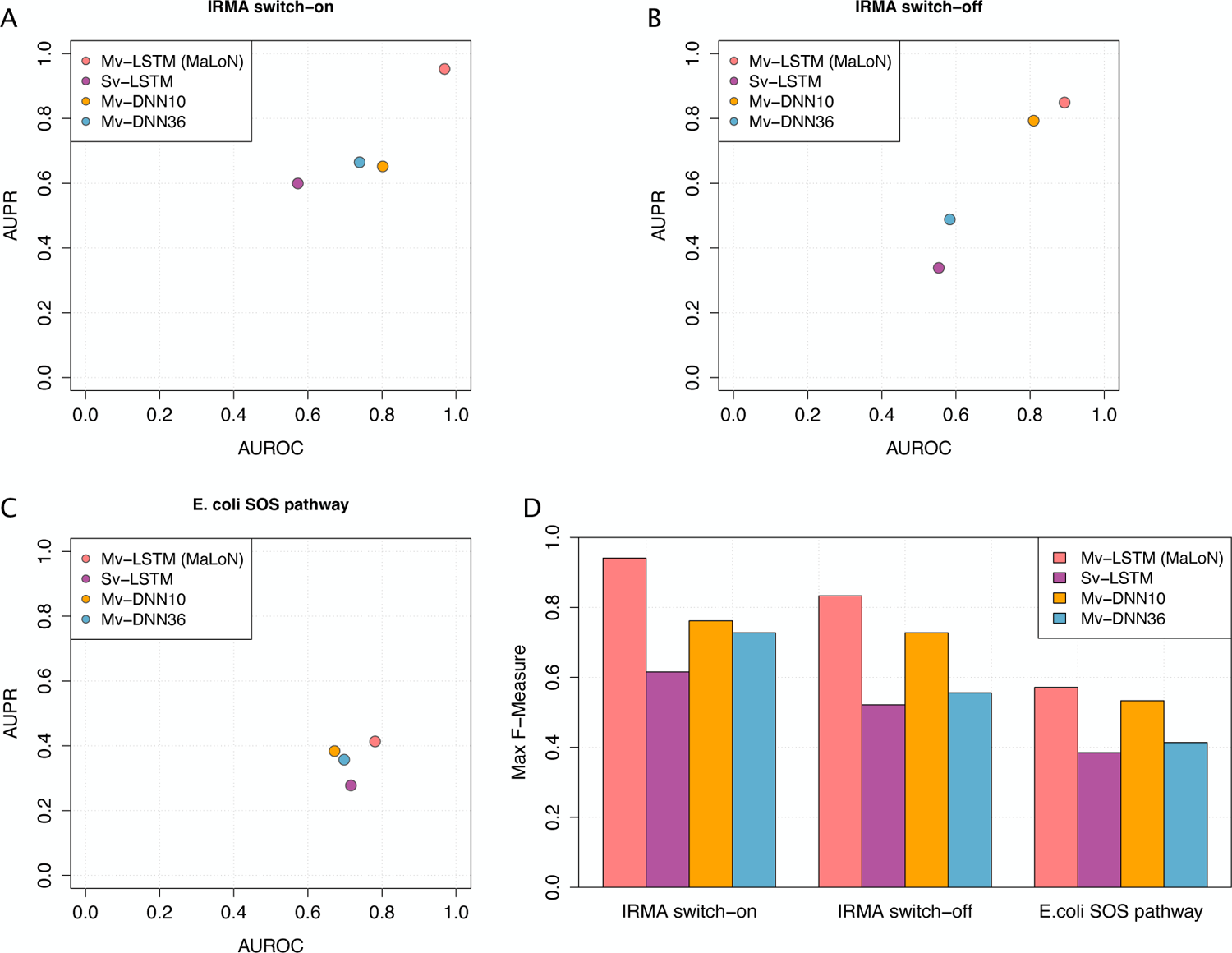
Comparison of MaLoN (Mv-LSTM) and ablated models. Sv-LSTM, single-view LSTM; Mv-DNN10, multi-view DNN with a hidden layer vector size of 10; Mv-DNN36, multi-view DNN with a hidden layer vector size of 36. (A-C) AUROC, AUPR, and (D) Max F-measure were used as metrics. Datasets used are indicated on the *x* axis in (D).

### Interpretation of the inferred dynamic regulation maps

We visualized the dynamic regulation map inferred for the datasets used in the evaluation. In the IRMA switch-on dataset with the focus on the inferred regulation of the GAL4, the regulation from CBF1 and from GAL80 had high scores at different times (Fig. 6A). Such a temporal switch in gene regulation in the IRMA dataset has not been reported. This finding might indicate that the transcriptional system in *S. cerevisiae* generates a specific order of gene expression. In the *E. coli* SOS Pathway dataset, we observed a delay in the regulation from the hub regulator *lexA* to each gene at specific periods (Fig. 6B). *UvrA* and *uvrD* function in nucleotide excision repair and *ruvA* functions in double-strand break repair; they repair DNA by various mechanisms, and may regulate other genes in response to perturbations [23].

**Fig. 6:**
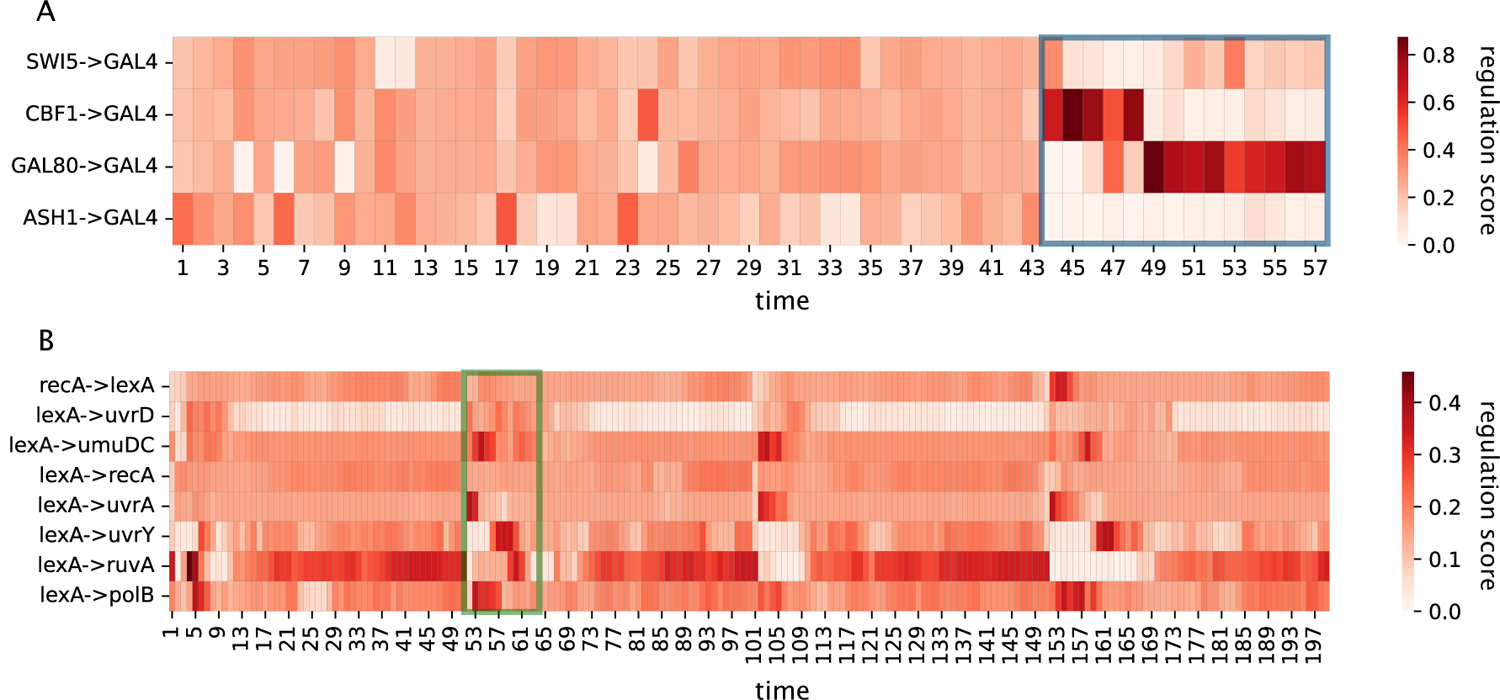
Visualization of inferred dynamic gene regulation maps for IRMA switch-on and *E. coli* SOS pathway datasets. (A) Map for regulations from other genes to GAL4 in the IRMA switch-on dataset. The high scores (blue box) at t=43-47 for the regulation from CBF1 to GAL4, and at t=48-56 for the regulation from GAL80 to GAL4. (B) Maps for true regulation in the *E. coli* SOS pathway dataset. At t = 52-63 (green box), regulation from the hub gene *lexA* was activated to *uvrA* and *uvrD* and then to *umuDC* and *polB*.

## Discussion

In this study, we have proposed a novel method MaLoN, which accurately infers gene regulatory networks from *in vivo* time-series gene expression data. MaLoN is an unsupervised method that combined LSTM with a multi-view attention mechanism. LSTM is a deep learning architecture that can effectively learn time-series data, and multi-view attention is a mechanism for regression of target gene expression that considers which regulators and time points are important. In MaLoN, the time-series expression of target genes is regressed from that of other genes, and then multi-view attention mechanism is used to infer not only which genes but also which time points are important for the regression. Therefore, this method enables inference of gene regulatory networks that can express as dynamic changes in gene regulation, which is not possible with the existing methods. Using MaLoN, we can effeciently infer gene regulatory networks from the *in vivo* data, which existing regression tree models cannot do. MaLoN is implemented to have the constraint that gene expressions at time *t* must be regressed only from the past. This cannot be achieved with architectures such as Transformer [25], which allows regression from any time.

The ablated models showed that both the multi-view attention mechanism and LSTM architecture improved performance through the suppression of false positives. To detect causality in gene regulation, it is necessary to capture changes in expression of a target gene at a limited number of time points and to search for genes with expression patterns that can explain the changes. In the regression tree-based methods, the tree is split at the time when the expression of the target gene and upstream genes consistently changes with time lags. The causality is then inferred by calculating a score based on which gene was used to split the tree. Therefore, to detect causality, these methods consider only local information in a time-series. Such local search may lead to false positives due to the lack of global features of expression dynamics necessary for causality detection. In contrast, MaLoN uses LSTM to detect causality by memorizing the global time-series characteristics, which is effective for narrowing down the candidate regulators and thus reducing false positives. The scoring at each time point by multi-view attention mechanism also contributes to the suppression of false positives by the detecting expression changes. The ablated models, showed that both the adoption of the multi-view attention mechanism and LSTM suppressed false positives, with the multi-view attention mechanism having an especially large effect. These results suggest that a model that can represent dynamic changes in gene regulation and directly learn the global time-series information can efficiently infer gene regulatory networks from time-series expression data.

A key feature of MaLoN is the ability to infer dynamic gene regulation from time-series gene expression alone. In this study, we inferred dynamic gene regulation in the *S. cerevisiae* IRMA switch-on and switch-off datasets and the *E. coli* SOS Pathway dataset. In the *E. coli* dataset, the hub *lexA* regulated other genes in a specific order (Fig. 6B). The *uvrA* and *uvrD* genes were activated early, followed by activation of *umuDC* and *polB*. The SOS system in *E. coli* is responsive to DNA damage [26]. The transcription of *uvr* is activated immediately after the SOS response, and then, upon continued DNA damage, the transcription of DNA repair DNA polymerase genes *polB*, *umuC*, and *umuD* is activated. This provides evidence that MaLoN can find the true regulatory relationships without prior knowledge. Our results are partially consistent with existing knowledge, but whether all of them are valid is unknown. Experimental validation of the inference by MaLoN may reveal unknown dynamic gene regulation.

Although the above issues remain, we expect MaLoN to have various applications in molecular cell biology; (i) it can be used to track the dynamics of cell state transitions by inferring dynamic gene regulation, i.e., at what time and with the involvement of which regulatory factors the transitions occur; (ii) the inferred dynamic gene regulatory networks will enable various network analyses focused on temporal behavior. Many previous studies have considered gene regulatory networks as a static Boolean networks and have analyzed them without considering the edge dynamics, such as identification of upstream factors that regulate the target genes or mapping information such as time-series expression of the node genes [27]. Several studies have modeled expression dynamics from inferred gene regulatory networks, but have not directly considered edge dynamics and require multi-omics data [28, 29]. In contrast, MaLoN can add information on the dynamics of edges only from transcriptome data and enable network analysis that can reveal the dynamic characteristics of both nodes and edges in terms of systems biology.

## Materials and Methods

### Multi-view attention LSTM model

MaLoN infers gene regulatory networks by learning regressions of gene expression using a LSTM model with multi-view attention mechanism for gene- and time-view and interpreting the attention map. Inference of a gene regulatory network is equal to constructing a directed network *G*(*V, E*) from time-series gene expression data **x**, where *V* is the set of nodes as genes and *E* is the set of edges as gene regulatory relationships. Edges from node *g_i_* to *g_j_*(*g_i_, g_j_ ∈ G*) are represented as *e_i,j_ ∈ E* and it means that gene *i* affects the expression of gene *j*. The gene expression data have *N* genes and *T* time points, and *x_i_*(*t*) is the expression of gene *i* at time *t*. MaLoN regresses target gene expression *x_i_*(*t*+1) from the other genes at the previous time points *x_−i_*(**t**)(**t** = *{*1, 2*, ·· ·, t}*), as represented by the following equation:

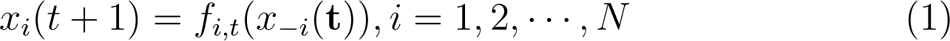

where *f_i,t_* is a nonlinear function defining the expression of a gene *i* at time *t* + 1 and this is learned independently for each of the *N* genes. The multi-view attention LSTM model was applied for representing *f_i,t_*. In this model, attention maps for the scoring of regulators to a target gene *i* are obtained as follows.

1. pre-training: the regression of target gene *i* at time *t* + 1 is learned from all other genes on the basis of equation (1).
2. Using the pre-trained model, the feature vector *h_−i,j_*(*t*) is calculated from *x_−i,j_*(*t*), which is the expression of gene *j* (the expression of all other genes is masked). This is repeated for all candidate regulators to obtain the feature matrix *H_−i_*(*t*) = *{h_−i,j_*(*t*)*|j* = 1, 2*,i −* 1*,i* + 1*, ···,N}*. The feature vector *h_−i_*(*t*) from all genes is also calculated.
3. The inner products of *H_−i_*(*t*) and *h_−i_*(*t*) are calculated as attention weights. This means quantifying which feature vectors obtained from which genes are similar to the vectors obtained from all genes. We assume that the feature vectors from all genes has enough information to regress the expression of the target gene, and thus genes similar to it have higher scores as regulator candidates for gene *i*.
4. Finally, *x_i_*(*t* + 1) is calculated through the attention layer and one fully connected layer. The attention weights calculated at each time point are obtained as an attention map (dynamic regulation map) of size (*T −* 1) *×* (*N −* 1) for a target gene *i*.

The regression of a target gene is learned to minimize the error between the predicted result and the expression data. We used the mean squared error as the loss function and stochastic gradient descent as the optimization method for all evaluations.

### Performance metrics

The Area Under the Receiver Operating Characteristic curve (AUROC) and Area Under Precision Recall curve (AUPR) were used as performance metrics; they are generally used in the evaluation of existing methods to infer gene regulatory networks [19]. AUROC is the area under the curve based on the true positive rate (TPR) and false positive rate (FPR). AUPR is the area under the curve based on precision and recall. TPR, FPR, precision, and recall are each represented as follows.

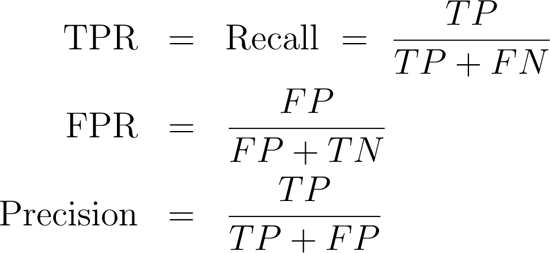

where *TP*, *TN*, *FP*, and *FN* are the numbers of true positives, true negatives, false positives, and false negatives, respectively. To obtain a unique gene regulatory network structure from the edge ranking, it is necessary to decide the confidence threshold and classify edges as positive or negative. To evaluate the ability to obtain a gene regulatory network structure that is similar to that of the true network, we used Max F-measure, which is F-measure as follows when the threshold that maximizes F-measure is adopted.

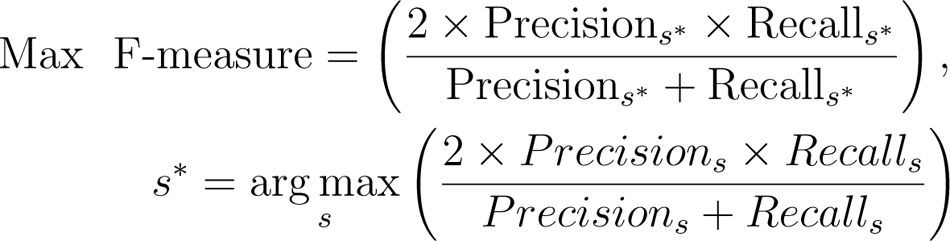

where Precision*_s_* and Recall*_s_* are obtained using a threshold *s*.

## Datasets

We used the *in vivo* time-series gene expression datasets obtained from *S. cerevisiae* and *E. coli*, for which true network structures are known. We inferred gene regulatory networks using these datasets and evaluated the performance with the metrics explained in the previous section. Information on each dataset is shown in Table 1.

### IRMA datasets

The *in vivo* reverse-engineering and modeling assessment (IRMA) dataset is a synthetic regulatory network of five genes in *S. cerevisiae* [22]. In this dataset, GAL4 and GAL80 interact when cultured cells are placed in a galactose medium (switch-on; 5 experiments), and stop to interact when in a glucose medium (switch-off; 4 experiments). The experiments were combined in each dataset.

### *Escherichia coli* SOS pathway dataset

The *E. coli* SOS pathway dataset is a regulatory network of 8 genes, which are activated after DNA damage. This network induces the expression of 30 genes that are associated with DNA damage tolerance and repair [23]. It contains time-series expression data from 4 experiments, which we conmbined.

## Code availablity

The source code of MaLoN is available from https://github.com/funalab/MaLoN.git

## Acknowledgements

We thank T. Kobayashi for helpful discussions on the difficulty of gene regulatory network inference. The research was funded by JST CREST Grant Number JPMJCR2011 including AIP Challenge program and JSPS KAKENHI Grant Numbers 21J20961 to Y.H. We are grateful to two professional editors from ELSS for careful editing of the manuscript.

## Author information Authors and Affiliations

Center for Biosciences and Informatics, Graduate School of Fundamental Science and Technology, Keio University, Yokohama, Kanagawa, 223-8522, Japan Yusuke Hiki, Department of Biosciences and Informatics, Keio University, Kanagawa, 223-8522, Japan, Yuta Tokuoka, Takahiro G. Yamada & Akira Funahashi

## Contributions

Y.H., T.G.Y and A.F. designed the conceptual idea and the study. Y.H. implemented the code of MaLoN, analyzed the results, and wrote the manuscript. Y.T. provided deep learning expertise and supervision. All authors read and approved the final manuscript.

## Ethics declarations Competing Interests

The authors declare that they have no competing financial interests.

