## Supplementary Material for "Inference of gene regulatory networks for overcoming low performance in real-world data"

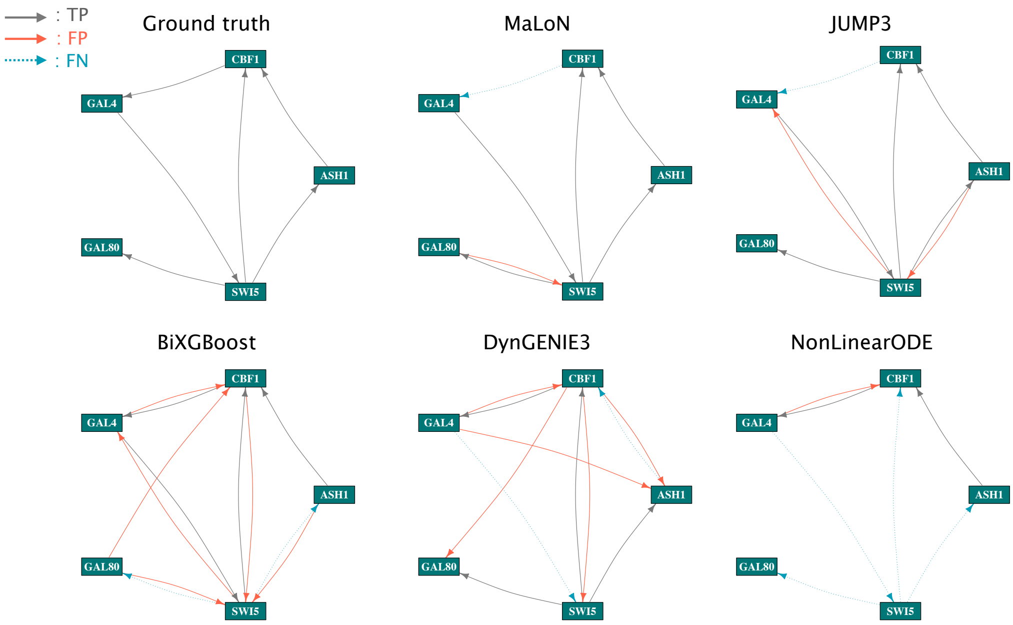


Supplementary Fig. 1: Visualization of the inferred gene regulatory network in the IRMA switch-off dataset. The nodes represent the 5 genes of this dataset. TP, true positive; FP, false positive; and FN, false negative. Ground truth is the true network, and the other networks were obtained by each method using the threshold with Max F-measure.

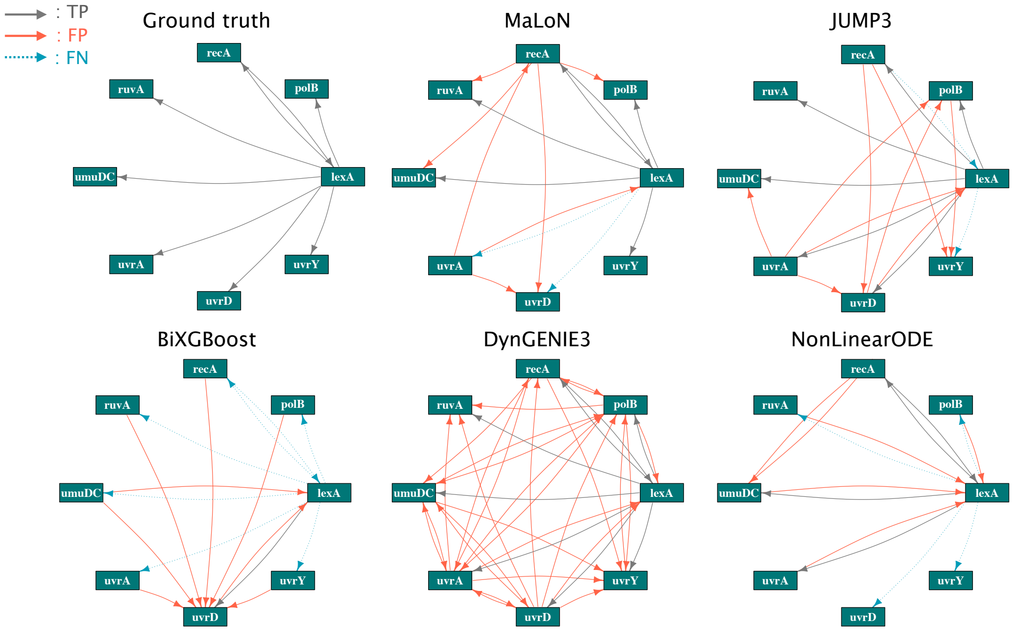


Supplementary Fig. 2: Visualization of the inferred gene regulatory network in the *E. coli* SOS pathway dataset. The nodes represent the 8 genes of this dataset. TP, true positive; FP, false positive; and FN, false negative. Ground truth is the true network, and the other networks were obtained by each method using the threshold with Max F-measure.

Supplementary Table 1: Quantification of Max F-measure, precision and recall for each ablated model

| | | Mv-LSTM | Sv-LSTM | Mv-DNN10 | Mv-DNN36 |
|---|---|---|---|---|---|
| IRMA switch-on | Max F-measure | 0.941 | 0.615 | 0.762 | 0.727 |
| | Precision | 0.889 | 0.800 | 0.615 | 0.571 |
| | Recall | 1.00 | 0.500 | 1.00 | 1.00 |
| IRMA switch-off | Max F-measure | 0.833 | 0.522 | 0.727 | 0.556 |
| | Precision | 0.833 | 0.353 | 0.800 | 0.417 |
| | Recall | 0.833 | 1.00 | 0.667 | 0.833 |
| *E. coli* SOS pathway | Max F-measure | 0.571 | 0.385 | 0.533 | 0.414 |
| | Precision | 0.462 | 0.278 | 0.571 | 0.286 |
| | Recall | 0.750 | 0.625 | 0.500 | 0.750 |
